# The fate of horizontally acquired genes: rapid initial turnover followed by long-term persistence

**DOI:** 10.1101/2025.08.05.668751

**Authors:** Swastik Mishra, Käthe Weit, Martin J. Lercher

## Abstract

A major driver of bacterial evolution is horizontal gene transfer (HGT), the acquisition of genes from other strains or species. Transfers between closely related taxa are more likely to succeed, while the pervasive deletion bias of bacterial genomes drives frequent turnover of horizontally acquired genes. However, whether the rate of gene loss after acquisition is constant across lineages or time remains unclear. Here, we analyze a comprehensive dataset of bacterial genomes to infer the frequency, distribution, and retention of inter-phylum HGT events. The retention of inter-phylum gene transfers is highly skewed, with only a small subset of bacterial genomes accounting for the majority of such events. Most transferred genes are lost rapidly. Those genes that survive the initial purging are retained over long periods, are biased toward functions such as transport and metabolism, and have larger numbers of protein-protein interactions.

## Introduction

Horizontal gene transfer (HGT) is a key driver of prokaryotic evolution (1, 2), even if its role in environmental adaptations may have been overestimated (3). Unlike vertical inheritance, HGT facilitates the exchange of genetic elements across phylogenetic boundaries, profoundly shaping bacterial evolution and ecological diversification.

Evidence indicates that most horizontally acquired genes originate from phylogenetically related donors rather than from distantly related taxa. This pattern likely reflects both selective pressures and mechanistic barriers to gene flow between divergent lineages, such as constraints imposed by genome architecture, functional compatibility, and integration with host regulatory networks (4–7). Although horizontal transfer of genes between bacteria from different phyla (inter-phylum HGT) is less frequent than within-phylum transfer, it has nonetheless been documented in notable cases, for instance, in the transfer of genes from archaea to bacteria at the origin of certain archaeal clades (8, 9). While such events are considered rare, their impact on recipient genomes can be substantial. Although it is difficult to estimate the frequency of inter-phylum HGT, Sheinman et al. (7) reported that approximately 8% of bacterial genomes harbor DNA segments identical to those in other phyla, indicating that inter-phylum gene transfer is not uncommon. Nevertheless, because these DNA segments may not always function as expressed genes, the functional relevance of such horizontally transferred sequences – and whether all bacterial clades are equally susceptible to inter-phylum HGT – remains unclear.

Furthermore, studies on sets of closely related species indicate that the retention of newly acquired genes is often transient; empirical studies indicate that the majority of horizontally transferred genes are rapidly lost following their initial acquisition (10–12). However, it is unclear if this pattern of rapid loss holds generally across bacteria.

Here, we employ a systematic, sequence-identity-based method for HGT inference to analyze patterns of gene loss across a comprehensive dataset of > 33,000 bacterial genomes. Our approach differentiates between very recent and slightly older inter-phylum HGT events and examines their persistence, functional categorization, and involvement in protein–protein interaction networks. The horizontal acquisition of potentially functional genes from other phyla is very rare; however, a quarter of the genomes that contain any inter-phylum gene acquisitions harbor genes from multiple gene families. We observe that horizontally acquired genes are initially lost at a high rate; those retained appear to be very stable afterwards and display distinct functional characteristics compared to genes eliminated early. Collectively, these findings support a two-phase model of post-transfer gene loss and broaden our understanding of the evolutionary forces shaping the retention of horizontally acquired genes.

## Results and Discussion

### Inference of recent inter-phylum gene transfers

We analyzed 35,439 gene families from 33,918 extant bacterial genomes, obtained from the EggNOG database v6 (13), which clusters genes into non-supervised orthologous groups (NOGs; referred to as gene families below). Horizontal gene transfer (HGT) was inferred by identifying, within a gene family, pairs of genes with high sequence identity that are found in two different phyla as defined by NCBI Taxonomy (14). In this framework, a horizontally acquired gene is represented as a pair of homologous genes from different taxa. Sequence identity — the percentage of identical amino acids – was calculated from the multiple sequence alignments (MSA) of EggNOG gene families. The distribution of such gene pairs across percent sequence identities is shown in Fig. 1. The direction of transfer (i.e., which gene in the pair is from the donor taxon and which from the recipient) was inferred by examining the outgroup of the most recent common ancestor in the gene tree; if one phylum is represented in the outgroup and the other is not, directionality can be assigned, i.e., we can identify which genome is the recipient and which is the donor (see Methods). The vast majority of gene families contain no inter-phylum gene pairs with sequence identities above 80%. We find candidate inter-phylum HGT gene pairs in 796 gene families and 4,445 genomes for downstream analyses.

**Fig. 1.**
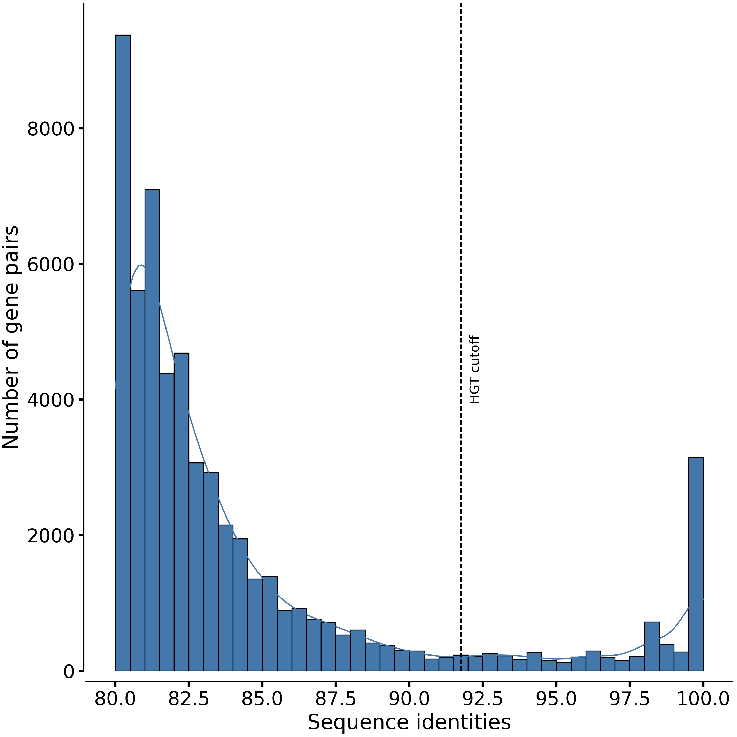
Recent inter-phylum HGTs can be distinguished from vertically inherited genes from sequence identities. Distribution of sequence identities of gene pairs in a gene family such that the pair of genes are from two different phyla. The dotted line indicates the estimated cutoff for HGT inference, above which we consider the gene pairs to be horizontally transferred.

The overall distribution of inter-phylum gene pairs exhibits three major trends: a prominent peak at 100% sequence identity, an approximately uniform distribution at intermediate sequence identities, and a gradual increase as sequence identity decreases further. For comparison, we performed the same analysis to identify candidate inter-class and inter-order HGT pairs. For these, the increase in frequency begins at much higher sequence identities (Suppl. Fig. S1). This pattern suggests that gene pairs with lower sequence identity primarily result from vertical inheritance, as vertically inherited genes within the same family will be more similar among more closely related taxa. We conclude that pairs of genes from different taxonomic groupings with high sequence identities, including those at 100% sequence identity, are predominantly the results of HGT. Results are very similar when considering only those gene pairs where both genes are from genomes with complete assemblies (Suppl. Fig. S4, i.e. assemblies in NCBI RefSeq that are labeled as “Complete Genome”; which are 1191 genomes, or 26.79% of the 4445 genomes involved in HGT in our dataset), arguing against any major role of contaminations and assembly artifacts in the inferred interphylum gene transfers.

Accordingly, we applied a sequence identity cutoff of 91.75% to identify gene pairs from inter-phylum HGT. This threshold corresponds to the point at which the smoothed frequency distribution of sequence identities flattens when moving from lower to higher sequence identities (see Methods and Suppl. Fig. S3).

For two highly similar homologous genes found in two genomes from different phyla, sequence identity is an approximate measure of the number of substitutions since their last common ancestor, and can thus serve as a proxy for the time that has passed since the HGT event. Even without the assumption of a molecular clock, one can distinguish approximately between recent and older transfers by considering the inter-phylum gene pairs that are identical versus the ones that are not. Accordingly, we classified pairs with 100% sequence identity as *recent* transfers, whereas those with sequence identities below 100% but above the cutoff are regarded as *older* (albeit still relatively recent) transfers. The majority of inter-phylum gene pairs identified as HGTs (84.51% or 4,245 of 5,023) are older transfers.

### Inferences of HGT from high sequence similarity between phyla are highly precise

Could inferred HGT pairs contain false positives, i.e. vertically inherited genes from highly conserved gene families? We estimated that the chance of finding even one gene pair in our recent or older transfer datasets that is actually due to vertical inheritance from the common ancestor of two phyla (the family-wise error rate; see Methods) is essentially zero. In addition to theoretical calculations, we performed an empirical survey of gene families in the dataset. Across all gene families examined, we did not observe any gene families that contained (i) at least one pair of genes from different phyla with > 80% amino acid sequence identity and (ii) monophyletic gene trees with respect to the represented phyla. This indicates that such highly similar cross-phyla pairs are not observed in typical vertically inherited gene families at deep taxonomic levels in our dataset, and supports the interpretation that the cross-phyla identities at the HGT cutoff we detect are unlikely to arise under a purely vertical null model. To further validate that inferred HGT pairs are statistical outliers compared to vertical inheritance, we computed z-scores for each transfer by comparing its sequence identity to the distribution of all non-transfer inter-phylum pairs within the same gene family and phylum pairing (see Methods). Both recent (*N* = 2,061, median *z* = 0.31) and older (*N* = 54,261, median *z* = 1.54) transfers showed significantly elevated sequence identity relative to the vertical background (Wilcoxon signed-rank test, *p* < 10^−74^ for both categories); see Suppl. Fig. S2. Older transfers exhibited even stronger deviations (Mann-Whitney *U* test, *p* < 10−245), suggesting that genes from highly conserved gene families are enriched in recent transfers, where they are quickly purged. Thus, the transfers inferred here are extreme outliers in sequence similarity and are unlikely to result from vertical inheritance.

### Inter-phylum gene acquisitions are concentrated in a few genomes

The distribution of the number of interphylum HGTs per genome is highly skewed, with most genomes (98.16%; 33294 of 33918) not hosting any interphylum HGTs that are recent enough to pass our filters. Only 0.57% (195 of 33918) of the genomes in our dataset (195 out of 33,918) are involved in any relatively recent inter-phylum HGT event for which we could infer the direction of transfer. Of these, only about a quarter (53 out of 195) show evidence of recent transfers, while most (180 out of 195) only harbour older transfers. Only about a quarter of the genomes involved in inter-phylum HGTs (26.7%, or 52 out of 195) received genes from more than one gene family, and only 3.1% (6 out of 195) received genes from more than 5 gene families (Fig. 2a) These genomes belong to diverse phyla, with no single phylum dominating the distribution of inter-phylum HGTs (see Suppl. Table S1 for lineage information). Fig. 2b shows that recent inter-phylum HGTs are also concentrated in a handful of bacterial genomes.

**Fig. 2.**
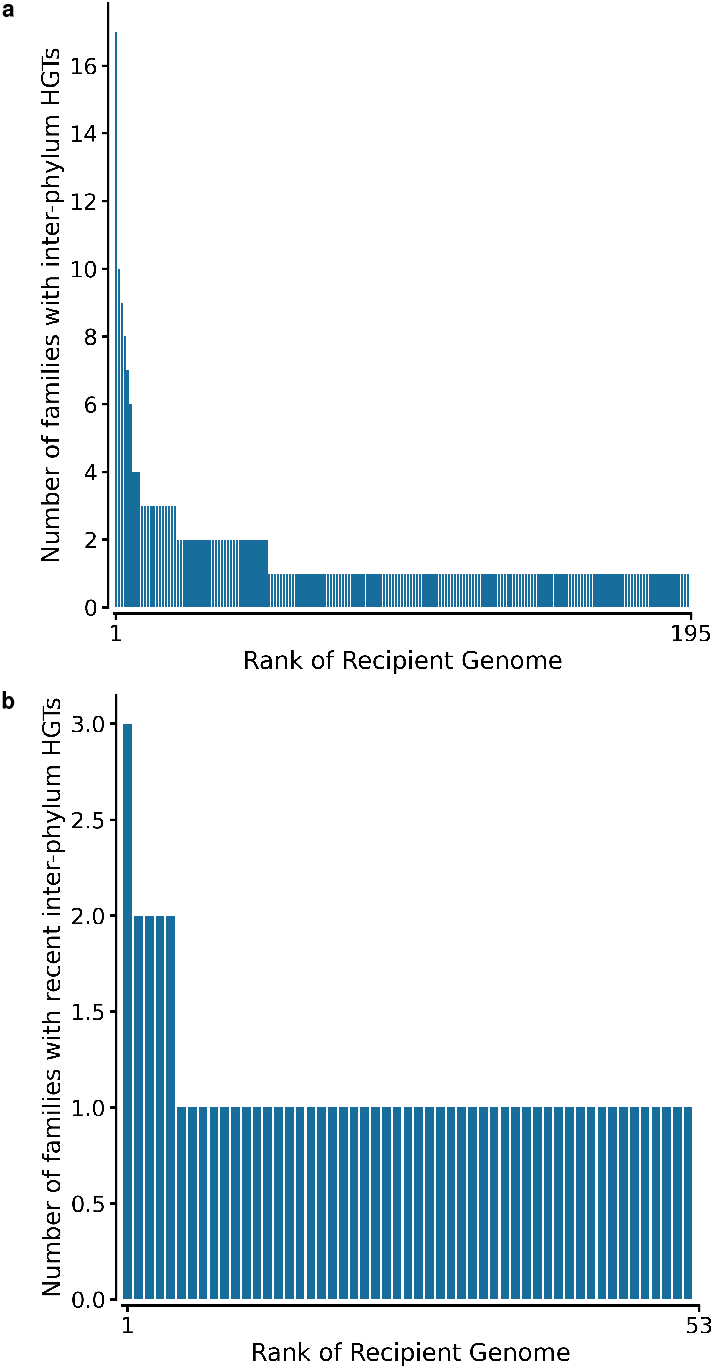
The distribution of inter-phylum HGTs is strongly skewed. Rank plot of the number of inter-phylum gene acquisitions per genome. The x-axis shows the genomes ranked by the number of inter-phylum HGTs, while the y-axis shows the number of gene families with inter-phylum HGTs where the direction of transfer is known. **(a)** All inter-phylum HGTs. **(b)** Recent inter-phylum HGTs.

### The loss rate of horizontally acquired genes shows two phases

The pronounced peak of inter-phylum gene pairs at 100% sequence identity in Fig. 1 indicates that a large proportion of horizontally acquired genes are lost shortly after their acquisition: only few of these genes are still present after enough time has passed for the gene pair to have accumulated even a small number of substitutions. The near-uniform distribution across lower identity levels implies that very few genes are lost at later stages. The pronounced peak at 100% sequence identity, immediately followed by a near-uniform distribution, is also seen for inter-order and interclass HGT (Suppl. Fig. S1). Thus, gene loss following HGT appears to generally follow a two-phase dynamic: an early phase of rapid loss, followed by long-term retention of the remaining genes.

### Functional biases are elevated among genes that survive initial loss

How do horizontally acquired genes that persist in their host genome differ from those eliminated in the early, high-turnover phase? To answer this question, we used Clusters of Orthologous Groups (COG) functional category annotations in EggNOG. We performed Fisher’s exact tests to check if the proportion of pairs in a given COG category differs between those that are inferred to be transferred across phyla and those not inferred to be transferred (Fig. 3a), and between recent and older transfers (Fig. 3b). After correcting for multiple hypothesis testing (15), we observed that most COG categories that are associated with transport and metabolism (categories I, E, G, H, C) as well as secondary metabolite biosynthesis (Q) are significantly depleted in inter-phylum HGTs compared to non-transfers. On the other hand, categories that are enriched in gene transfers across phyla include those associated with (L) replication, recombination and repair, (J) translation, ribosomal structure and biogenesis, (V) defense mechanisms, (D) cell cycle control, cell division, chromosome partitioning, (K) transcription, (U) intracellular trafficking, secretion, and vesicular transport, and (P) inorganic ion transport and metabolism. This result is consistent with the complexity hypothesis (16), which posits that genes that are successfully transferred belong more frequently to operational functional categories, such as metabolism and transport, than to informational ones, such as transcription and translation.

**Fig. 3.**
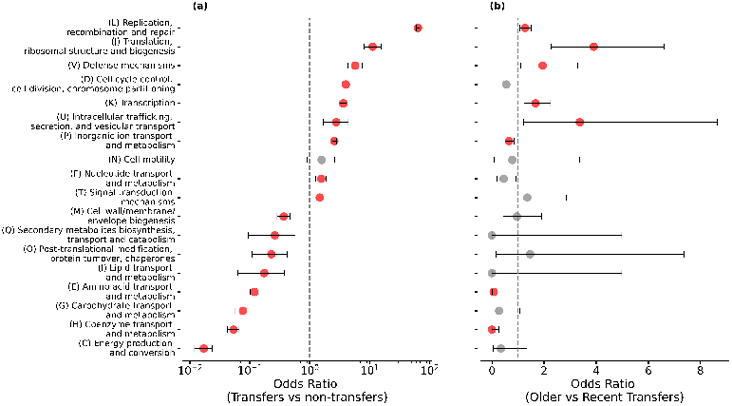
Recent and older inter-phylum transfers differ systematically in their functions, relative to non-transfer gene pairs. Odds ratios (OR) of COG categories enriched (OR>1) and depleted (OR<1) in (left) inter-phylum gene pairs inferred as transfers, vs non-transfers (right) older vs recent transfers. Error bars show 95% confidence intervals. Red markers indicate COG categories that are significantly enriched/depleted (adjusted *p* ≦ 0.05 after correcting for multiple testing). The dashed line at OR=1 indicates no difference between the categories.

Do genes that are retained after the initial phase of loss differ functionally from those that were gained recently? Except for inorganic ion transport and metabolism (P), we find that COG categories that are significantly depleted in older compared to recent transfers (Fig. 3b) are also significantly depleted in inter-phylum transfers overall (Fig. 3a). Conversely, COG categories that are significantly enriched in older versus recent transfers are either also significantly enriched in inter-phylum transfers overall or show no significant difference. This indicates that the functional biases observed among inter-phylum HGTs are amplified when considering only those genes that persist after the initial loss phase.

The rationale behind the complexity hypothesis is that genes in the information-processing functional categories are more complex in the sense of having more interaction partners, and are thus less likely to integrate successfully into the interaction network of a new host (16, 17). One therefore expects that genes transferred more frequently have a lower connectivity in terms of protein-protein interactions (PPI), a prediction that was previously confirmed in a much more limited dataset of older transfers (17).

We used the STRING database v11 (18) to obtain the number of PPI for every gene in our HGT dataset based on STRING’s own gene family clustering and a minimum STRING interaction score of 900 (max: 1000). In line with the Complexity Hypothesis, the mean PPI of inter-phylum gene pairs inferred as HGT, is an order of magnitude lower than that of non-transfer gene pairs (6.81 vs. 38.37, *p* < 1 × 10^−5^ from a permutation test). Although we observe a marginal negative correlation between the number of HGTs inferred for a gene family and the mean PPI of the gene family (Spearman’s *ρ* = -0.14 and -0.11 for recent and older transfers, respectively), neither of these correlations is statistically significant (*p* = 0.08 and *p* = 0.18 respectively; see Suppl. Fig. S5).

However, we find that the mean PPI across all gene pairs involved in older transfers is significantly higher than that across all gene pairs in recent transfers (6.95 vs. 5.99, *p* = 0.033 from a permutation test). Thus, genes involved in older transfers have on average one more protein-protein interaction partner than those involved in recent transfers.

## Discussion

Our analysis shows that inter-phylum transfers of potentially functional genes are rare: less than 2% of genomes have been involved – either as donor or recipient – in any inter-phylum HGTs that our methodology can infer. However, when focusing on those events for which we can infer the direction of transfer, a quarter of those genomes that contain any interphylum gene acquisition harbor acquisitions from multiple gene families. We observed that the probability of a randomly selected genome harboring an inter-phylum gene acquisition is 0.57%. Consequently, if all inter-phylum gene acquisitions occurred independently, the expected probability for a genome to acquire a second acquisition after the first would again be 0.57%, two orders of magnitude lower than the observed 26.7%. This observation suggests that once a genome has acquired a gene from another phylum, it becomes considerably more likely to acquire additional inter-phylum genes.

When inter-phylum gene acquisitions occur, the acquired genes follow a biphasic trajectory. A brief, intense purge removes most of the newcomers, while the remaining genes appear to be surprisingly stable and are retained over extended time periods. The fast, early attrition matches qualitatively the findings of Puigbò et al. (10), who reported a loss rate roughly three times higher than the rate of acquisition. Lineage-specific bursts of inter-phylum gene acquisitions have been described for individual species and genera (19, 20), lending further support to our findings.

The two phases of losing genes acquired through interphylum HGT likely reflect the same evolutionary processes that are responsible for the pronounced differences between intra-population polymorphisms, which are characterized by negative selection and drift, and inter-population substitutions, characterized by positive selection (21, 22). Thus, it appears likely that the most recent inter-phylum HGTs detected in our study are not fixed in the respective populations, but are polymorphism specific to the genomes that were deposited in the EggNOG database.

The complexity hypothesis (16) predicts that operational genes are more frequently transferred than informational ones. Our results align with this prediction: compared to very recent inter-phylum gene acquistions, older (yet still relatively recent) transfers are depleted in categories such as transcription, translation, replication/recombination/repair, intracellular trafficking, and defense – several of which represent informational functions. In contrast, genes that survived the early loss phase predominantly participate in operational roles, such as transport and metabolism.

The original formulation of the complexity hypothesis emphasized – without empirical support – the greater connectivity and functional entanglement of informational genes, a view that was supported by a more limited, earlier analysis of older transfers across different phylogenetic distances (17). However, our data reveal a different trend. Genes that undergo initial purging of horizontally acquired genes have, on average, more interaction partners than the most recent transfers. Thus, the observed functional biases likely reflect functional adaptations more than the sheer complexity initially proposed.

Our sequence similarity-based HGT inference approach offers precision that – based on neighboring co-acquisitions – is comparable to the best state-of-the-art methods at highest stringency, but focussed on relatively recent HGT events. However, it is inherently limited to detecting HGT between gene pairs covered by the dataset examined. Increasing both genome and gene family diversity would enable more accurate detection of gene similarities across clades, yielding a more complete picture of inter-phylum HGT. Sheinman et al. (7) observed that 8% of bacterial genomes share identical DNA segments with other phyla. That we observed a lower number of less than 2% likely reflects our focus on genes rather than DNA segments, as well as a more limited dataset. While the results of the two studies are not directly comparable, this discrepancy suggests that most inter-phylum DNA segment exchanges may not yield functional genes.

Together, our findings clarify the rarity and rapid attrition of inter-phylum horizontal gene transfers in bacteria, while highlighting the selective processes shaping their long-term retention. Future studies that combine broader genome sampling will be essential to map the ecological and molecular contexts in which such transfers persist.

## Materials and Methods

### Data and implicit phylogenetic method for HGT inference

Sequence identity, the number of substitutions, mismatches, and other relevant metrics were calculated for every gene pair within each NOG using multiple sequence alignments (MSA) from the EggNOG database. EggNOG also provides COG functional annotations for each gene family. Taxonomic information was retrieved from the NCBI Taxonomy via the Entrez API implemented through the Biopython library (14, 23). To infer the direction of HGT, gene trees available in EggNOG were used to identify appropriate out-groups. PPI data were obtained from STRING v11, considering only pairs of gene families with a minimum interaction score of 900 out of 1000 (18). Chromosomal positions of genes were also extracted from the STRING database.

To improve tractability and reduce noise, analyses were performed on a curated subset of the data. Specifically, only NOGs containing at least 10 taxa (to exclude very small groups) and a maximum of 2,000 genes were included. This subset comprised 33,918 genomes representing 35,439 NOGs across all bacterial phyla. For downstream analyses in this study, we further focused only on the 4,445 genomes containing 796 NOGs in which gene pairs exhibited sequence identity above 80% and originated from two different phyla. A subset of these gene pairs are inferred to be horizontally transferred based on our sequence identity cutoff.

Although recent transfers are defined as those with 100% sequence identity, to consider older transfers we identify a sequence identity cutoff based on the point where distribution of sequence identities flattens out. The distribution was smoothed using a rolling average with a window size of 5% sequence identity across bins of width 0.5%, excluding the bin for identical sequence matches. The cutoff was set at the point where the derivative of the smoothed distribution reached zero, indicating a plateau in the distribution.

### Family-wise error rate for recent HGT

We assume that substitutions at each site follow a Poisson process. Then, the probability that a site undergoes no amino acid substitutions over a branch length *d* (the mean number of substitutions per site) is *e*^−*d*^. For a gene of length *L*, the probability that all sites are unchanged is then

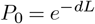

Across *M* independent gene pairs, the probability that no pair is identical by chance is (1 −*P*_0_)^*M*^. Then, the probability that at least one gene pair is identical by chance – the family-wise error rate (FWER) – is one minus this probability, i.e.,

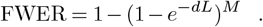

For our dataset, we estimate *d* as 4.94 substitutions per site, *M* as ≈ 10^9^ (number of gene pairs), and *L* as 267.41 (average gene length in number of amino acids). This leads to an estimate of FWER ≈ 0, i.e., a very low probability of false positives in our HGT inference of gene pairs with 100% sequence identity across phyla.

A similar calculation can be performed for any sequence identity cutoff *α*, for the FWER of older transfers. For a gene of length *L*, the number of unchanged sites follows K ∼ Binomial(*L, P*_0_). The probability of at least *α* sequence identity is

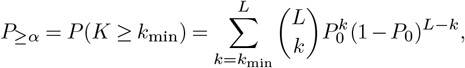

where *k*_min_ = ⌈*αL*⌉ is the minimum number of unchanged sites required to achieve a maximum fraction *α* identity (i.e. for a cutoff of 91.75%, *α* = 0.9175). To calculate *P*_≥*α*_, we can therefore calculate the survival function (i.e. 1-CDF) of the binomial distribution with parameters L and *p*_0_. Across *M* independent gene pairs, the family-wise error rate is

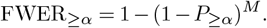

Using the values of *d, L*, and *M* as before, we obtain FWER_≥0.9175_ ≈ 0, indicating negligible false positive risk under this null for inter-phylum transfers.

### Z-score validation of HGT outliers

For each inferred HGT pair between two phyla in a gene family, we calculated a z-score quantifying deviation from vertical inheritance. We extracted all pairwise sequence identities between the same two phyla in that family, excluding inferred HGT pairs, and computed 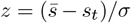, where *s*_*t*_ is the transfer’s sequence identity, 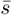 is the mean background identity, and *σ* is the standard deviation. Positive z-scores indicate transfers with higher similarity than the vertical inheritance background. Pairs with fewer than two background comparisons in the gene family or with zero standard deviation were excluded. We tested deviation from zero using the one-sample Wilcoxon signed-rank test (one-tailed) and compared recent versus older transfers using the Mann–Whitney U test (two-tailed).

### Outgroup for inferring direction of transfer and co– transfers

For any potential HGT in a gene family, the pair of genes must have two different taxonomic groupings at the taxonomic level of interest. Let A and B be two taxonomic groupings represented in a gene family tree, with gene members *A*_1_ and *B*_1_. If the outgroup of *A*_1_ and *B*_1_ has genes from A and not from B, then the direction of transfer is from the taxon containing *A*_1_ to the taxon containing *B*_1_, and vice versa. If two pairs of genes are inferred to be transferred from the same donor taxon to the same recipient taxon, we define it as a co-transfer. If they have the same recipient taxon, but not necessarily the same donor taxon, we define it as a co-acquisition. We limit our analysis of co-transfers and co-acquisitions to the gene pairs where the direction of transfer could be inferred.

### Functional categories for PPI analysis

We use the COG (Clusters of Orthologous Groups) functional categories as defined in NCBI COG (24), since EggNOG gene families already use these annotations for categorizing the NOGs. Fisher’s exact test was performed with a contingency table comparing the number of recent vs older HGT gene pairs, in a COG vs those not in the COG. The reliance on gene pairs introduces a potential for overestimation: in cases where several genomes within the same family are highly similar and only one belongs to a different phylum, multiple gene pairs may be labeled as HGT even if they result from a single transfer event.

Permutation tests were performed with 104 iterations of randomly shuffling whether a gene pair is a recent or older transfer, and then calculating the mean PPI for the pair of genes. The p-value is calculated as the fraction of iterations where the mean PPI of the older transfers is greater than that of the recent transfers. These gene families across the two sets are not independent, since a gene family can have both recent and older transfers. However, the permutation test is still valid, since the null hypothesis is that the mean PPI of the two sets of gene pairs is equal.

## Code and data availability

The code used for the analyses in this study is available on Gitlab (https://gitlab.cs.uni-duesseldorf.de/general/ccb/imli).

The corresponding data is available at Zenodo (https://doi.org/10.5281/zenodo.16745487).

## Competing interests

The authors have declared no competing interest.

## Funding

This work was supported by the Deutsche Forschungsgemeinschaft (DFG, German Research Foundation) through CRC 1310 and by the Volkswagenstiftung through the “Life?” initiative.

## Notes

https://gitlab.cs.uni-duesseldorf.de/general/ccb/imli

### Summary of Updates

added several tests to confirm inferred HGT events are unlikely to be false-positives/vertically inherited genes; enrichment analysis to compare HGT to non-HGT genes; verified main results with subset containing only complete genomes, excluding low quality genome assemblies.

## Bibliography

1. Brian J. Arnold, I.-Ting Huang, and William P. Hanage. Horizontal gene transfer and adaptive evolution in bacteria. Nature Reviews Microbiology, 20(4):206–218, April 2022. ISSN 1740-1534. doi: 10.1038/s41579-021-00650-4.

2. Csaba Pál, Balázs Papp, and Martin J. Lercher. Adaptive evolution of bacterial metabolic networks by horizontal gene transfer. Nature Genetics, 37(12):1372–1375, December 2005. ISSN 1546-1718. doi: 10.1038/ng1686.

3. Swastik Mishra and Martin J Lercher. Streamlined genomes, not horizontal gene transfer, mark bacterial transitions to unfamiliar environments | bioRxiv. bioRxiv, 2024, 2024. doi: 10.1101/2024.12.27.630308.

4. Christina L Burch, Artur Romanchuk, Michael Kelly, Yingfang Wu, and Corbin D Jones. Empirical Evidence That Complexity Limits Horizontal Gene Transfer. Genome Biology and Evolution, 15(6):evad089, May 2023. ISSN 1759-6653. doi: 10.1093/gbe/evad089.

5. Robert G. Beiko, Timothy J. Harlow, and Mark A. Ragan. Highways of gene sharing in prokaryotes. Proceedings of the National Academy of Sciences of the United States of America, 102(40):14332–14337, October 2005. ISSN 0027-8424. doi: 10.1073/pnas.0504068102.

6. Heather L. Hendrickson, Dominique Barbeau, Robin Ceschin, and Jeffrey G. Lawrence. Chromosome architecture constrains horizontal gene transfer in bacteria. PLoS Genetics, 14(5):e1007421, May 2018. ISSN 1553-7390. doi: 10.1371/journal.pgen.1007421.

7. Michael Sheinman, Ksenia Arkhipova, Peter F Arndt, Bas E Dutilh, Rutger Hermsen, and Florian Massip. Identical sequences found in distant genomes reveal frequent horizontal transfer across the bacterial domain. eLife, 10:e62719, June 2021. ISSN 2050-084X. doi: 10.7554/eLife.62719.

8. William F. Martin and Filipa L. Sousa. Early Microbial Evolution: The Age of Anaerobes. Cold Spring Harbor Perspectives in Biology, 8(2):a018127, December 2015. ISSN 1943-0264. doi: 10.1101/cshperspect.a018127.

9. Shijulal Nelson-Sathi, Filipa L. Sousa, Mayo Roettger, Nabor Lozada-Chávez, Thorsten Thiergart, Arnold Janssen, David Bryant, Giddy Landan, Peter Schönheit, Bettina Siebers, James O. McInerney, and William F. Martin. Origins of major archaeal clades correspond to gene acquisitions from bacteria. Nature, 517(7532):77–80, January 2015. ISSN 1476-4687. doi: 10.1038/nature13805.

10. Pere Puigbò, Alexander E. Lobkovsky, David M. Kristensen, Yuri I. Wolf, and Eugene V. Koonin. Genomes in turmoil: Quantification of genome dynamics in prokaryote supergenomes. BMC Biology, 12(1), December 2014. ISSN 1741-7007. doi: 10.1186/s12915-014-0066-4.

11. Weilong Hao and G. Brian Golding. The fate of laterally transferred genes: Life in the fast lane to adaptation or death. Genome Research, 16(5):636–643, January 2006. ISSN 1088-9051, 1549-5469. doi: 10.1101/gr.4746406.

12. Emmanuelle Lerat, Vincent Daubin, Howard Ochman, and Nancy A. Moran. Evolutionary Origins of Genomic Repertoires in Bacteria. PLOS Biology, 3(5):e130, April 2005. ISSN 1545-7885. doi: 10.1371/journal.pbio.0030130.

13. Ana Hernández-Plaza, Damian Szklarczyk, Jorge Botas, Carlos P Cantalapiedra, Joaquín Giner-Lamia, Daniel R Mende, Rebecca Kirsch, Thomas Rattei, Ivica Letunic, Lars J Jensen, Peer Bork, Christian von Mering, and Jaime Huerta-Cepas. eggNOG 6.0: Enabling comparative genomics across 12 535 organisms. Nucleic Acids Research, 51(D1): D389–D394, January 2023. ISSN 0305-1048. doi: 10.1093/nar/gkac1022.

14. Conrad L Schoch, Stacy Ciufo, Mikhail Domrachev, Carol L Hotton, Sivakumar Kannan, Rogneda Khovanskaya, Detlef Leipe, Richard Mcveigh, Kathleen O’Neill, Barbara Robbertse, Shobha Sharma, Vladimir Soussov, John P Sullivan, Lu Sun, Seán Turner, and Ilene Karsch-Mizrachi. NCBI Taxonomy: A comprehensive update on curation, resources and tools. Database, 2020:baaa062, January 2020. ISSN 1758-0463. doi: 10.1093/database/baaa062.

15. Yoav Benjamini and Yosef Hochberg. Controlling the False Discovery Rate: A Practical and Powerful Approach to Multiple Testing. Journal of the Royal Statistical Society: Series B (Methodological), 57(1):289–300, January 1995. ISSN 0035-9246. doi: 10.1111/j.2517-6161.1995.tb02031.x.

16. Ravi Jain, Maria C. Rivera, and James A. Lake. Horizontal gene transfer among genomes: The complexity hypothesis. Proceedings of the National Academy of Sciences, 96(7):3801– 3806, March 1999. ISSN 0027-8424, 1091-6490. doi: 10.1073/pnas.96.7.3801.

17. Ofir Cohen, Uri Gophna, and Tal Pupko. The Complexity Hypothesis Revisited: Connectivity Rather Than Function Constitutes a Barrier to Horizontal Gene Transfer. Molecular Biology and Evolution, 28(4):1481–1489, April 2011. ISSN 0737-4038. doi: 10.1093/molbev/msq333.

18. Christian von Mering, Lars J. Jensen, Berend Snel, Sean D. Hooper, Markus Krupp, Mathilde Foglierini, Nelly Jouffre, Martijn A. Huynen, and Peer Bork. STRING: Known and predicted protein–protein associations, integrated and transferred across organisms. Nucleic Acids Research, 33(Database Issue):D433–D437, January 2005. ISSN 0305-1048. doi: 10.1093/nar/gki005.

19. Hyaekang Kim, Woori Kwak, Sook Hee Yoon, Dae-Kyung Kang, and Heebal Kim. Horizontal gene transfer of Chlamydia: Novel insights from tree reconciliation. PLOS ONE, 13(4): e0195139, April 2018. ISSN 1932-6203. doi: 10.1371/journal.pone.0195139.

20. Alejandro Caro-Quintero and Konstantinos T Konstantinidis. Inter-phylum HGT has shaped the metabolism of many mesophilic and anaerobic bacteria. The ISME Journal, 9(4):958– 967, April 2015. ISSN 1751-7362. doi: 10.1038/ismej.2014.193.

21. Jun Gojobori, Hua Tang, Joshua M. Akey, and Chung-I Wu. Adaptive evolution in humans revealed by the negative correlation between the polymorphism and fixation phases of evolution. Proceedings of the National Academy of Sciences, 104(10):3907–3912, March 2007. doi: 10.1073/pnas.0605565104.

22. Austin L. Hughes, Robert Friedman, Pierre Rivailler, and Jeffrey O. French. Synonymous and Nonsynonymous Polymorphisms versus Divergences in Bacterial Genomes. Molecular Biology and Evolution, 25(10):2199–2209, October 2008. ISSN 0737-4038. doi: 10.1093/molbev/msn166.

23. Peter J. A. Cock, Tiago Antao, Jeffrey T. Chang, Brad A. Chapman, Cymon J. Cox, Andrew Dalke, Iddo Friedberg, Thomas Hamelryck, Frank Kauff, Bartek Wilczynski, and Michiel J. L. de Hoon. Biopython: Freely available Python tools for computational molecular biology and bioinformatics. Bioinformatics (Oxford, England), 25(11):1422–1423, June 2009. ISSN 1367-4811. doi: 10.1093/bioinformatics/btp163.

24. Michael Y. Galperin, Yuri I. Wolf, Kira S. Makarova, Roberto Vera Alvarez, David Landsman, and Eugene V. Koonin. COG database update: Focus on microbial diversity, model organisms, and widespread pathogens. Nucleic Acids Research, 49(D1):D274–D281, January 2021. ISSN 1362-4962. doi: 10.1093/nar/gkaa1018.

