## Supplementary Information for "The fate of horizontally acquired genes: rapid initial turnover followed by long-term persistence"

| Recipient Taxon ID | Phylum | Class | Order | Family | Genus | Species/Strain |
|---|---|---|---|---|---|---|
| 1120933 | Actinobacteria | Actinomycetia | Actinomycetales | Actinomycetaceae | Actinotignum | Actinotignum urinale DSM 15805 |
| 330214 | Nitrospirae | Nitrospira | Nitrospirales | Nitrospiraceae | Nitrospira | Nitrospira defluvii |
| 469615 | Fusobacteria | Fusobacteriia | Fusobacteriales | Fusobacteriaceae | Fusobacterium | Fusobacterium gonidiaformans ATCC 25563 |
| 552811 | Chloroflexi | Dehalococcoidia | - | - | Dehalogenimonas | Dehalogenimonas lykanthroporepellens BL-DC-9 |
| 653733 | Chrysiogenetes | Chrysiogenetes | Chrysiogenales | Chrysiogenaceae | Desulfurispirillum | Desulfurispirillum indicum S5 |
| 938289 | Firmicutes | Clostridia | Eubacteriales | - | Levyella | Levyella massiliensis |

**Table S1.** Lineage information for the genomes receiving more than 5 gene families with inter-phylum HGTs. If strain information was missing, the species name is shown instead.

### Supplementary Information

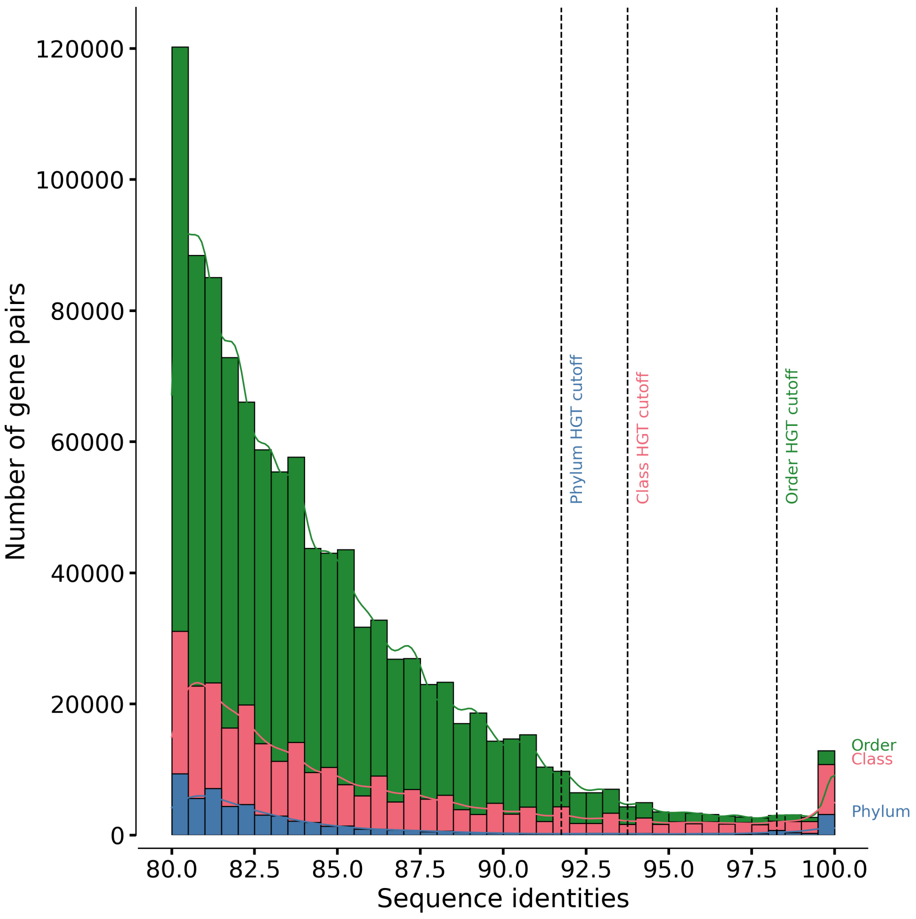


**Fig. S1. Distributions of the number of highly similar inter-clade gene pairs at different sequence identities are qualitatively similar at inter-phylum, inter-class, and inter-order levels.** The dashed lines indicate the estimated cutoff for HGT inference, above which we consider the gene pairs to be horizontally transferred.

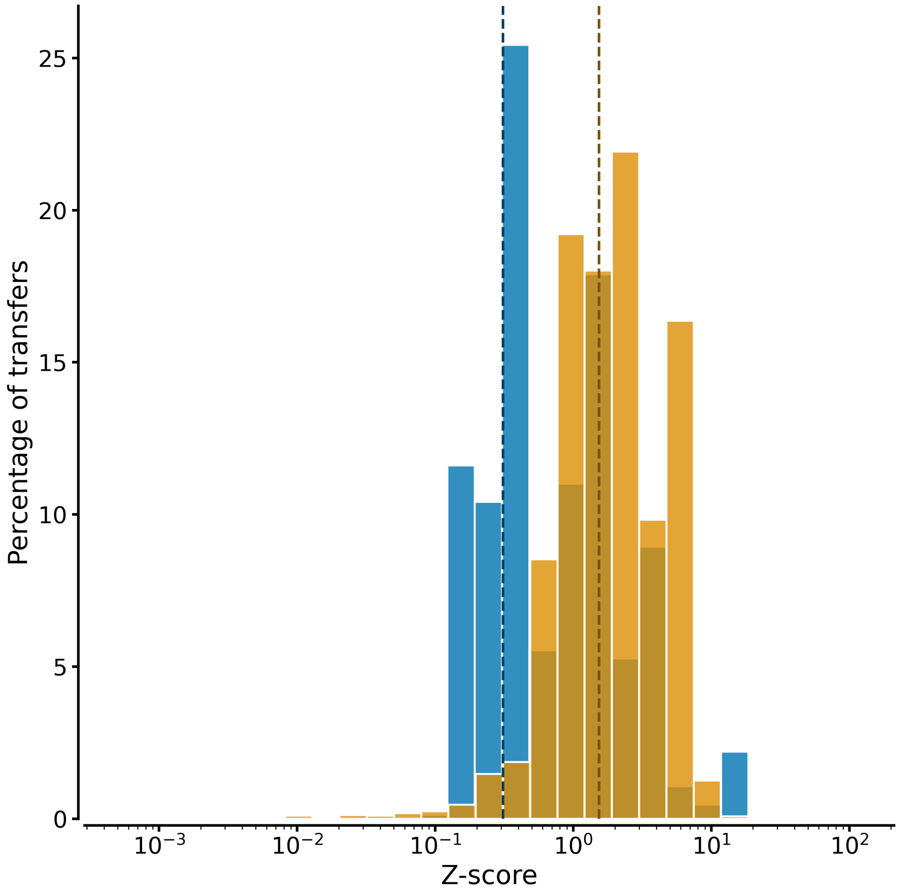


**Fig. S2. Z-score distributions show inter-phylum HGT pairs are outliers.** Recent (blue) and older (orange) transfers show significantly elevated sequence similarity compared to vertical inheritance backgrounds. Dashed vertical lines indicate median z-scores for recent ($Z = 0.31$) and older ($Z = 1.54$) transfers. Wilcoxon signed-rank test, $p < 10^{-74}$ for both categories.

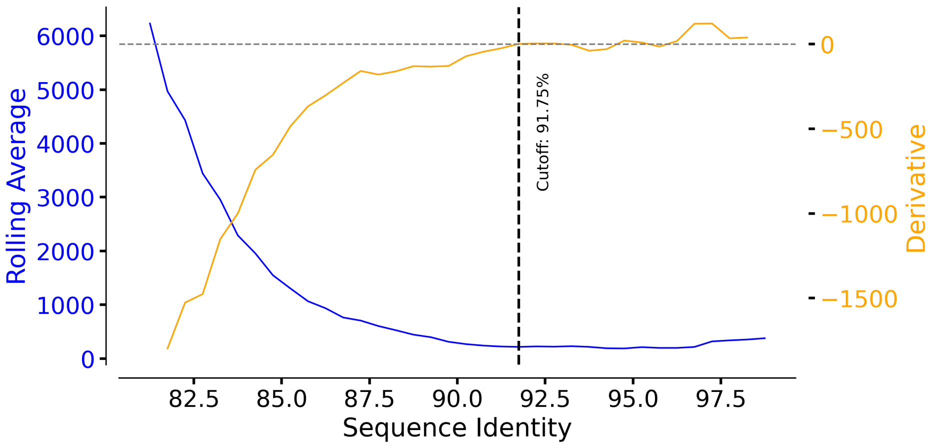


**Fig. S3. The cutoff for purely horizontal gene acquisition can be determined from the derivative of the sequence identity histogram.** The figure shows the rolling window average of the sequence identity histogram shown in Fig. 1 and its derivative. The dashed line indicates the cutoff for HGT inference, above which we consider the gene pairs to be horizontally transferred. The cutoff is chosen as the point where the rolling average flattens out, i.e., the derivative reaches zero for the first time when moving from lower to higher sequence identities.

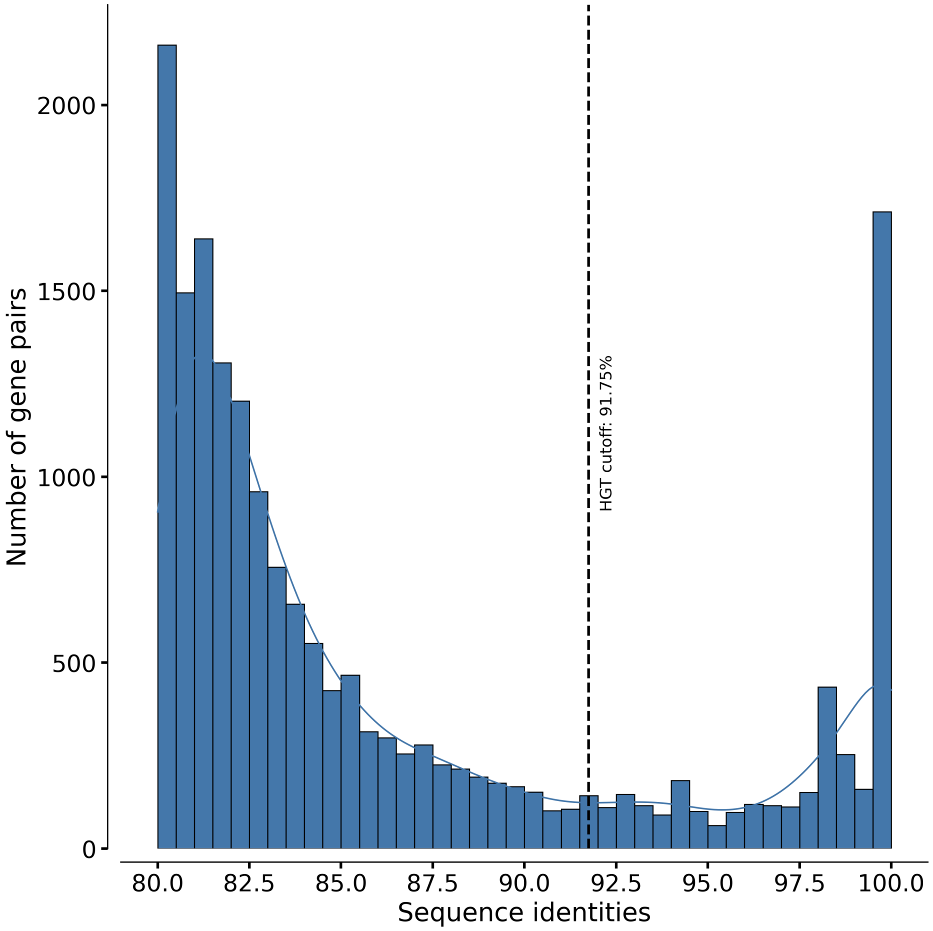


**Fig. S4. Distribution of sequence identities of gene pairs from different phyla in complete genomes.** Distribution of sequence identities of gene pairs in a gene family such that the pair of genes are from two different phyla, considering only genomes tagged as "Complete Genome" in the NCBI database. The dotted line indicates the estimated cutoff for HGT inference, above which we consider the gene pairs to be horizontally transferred.

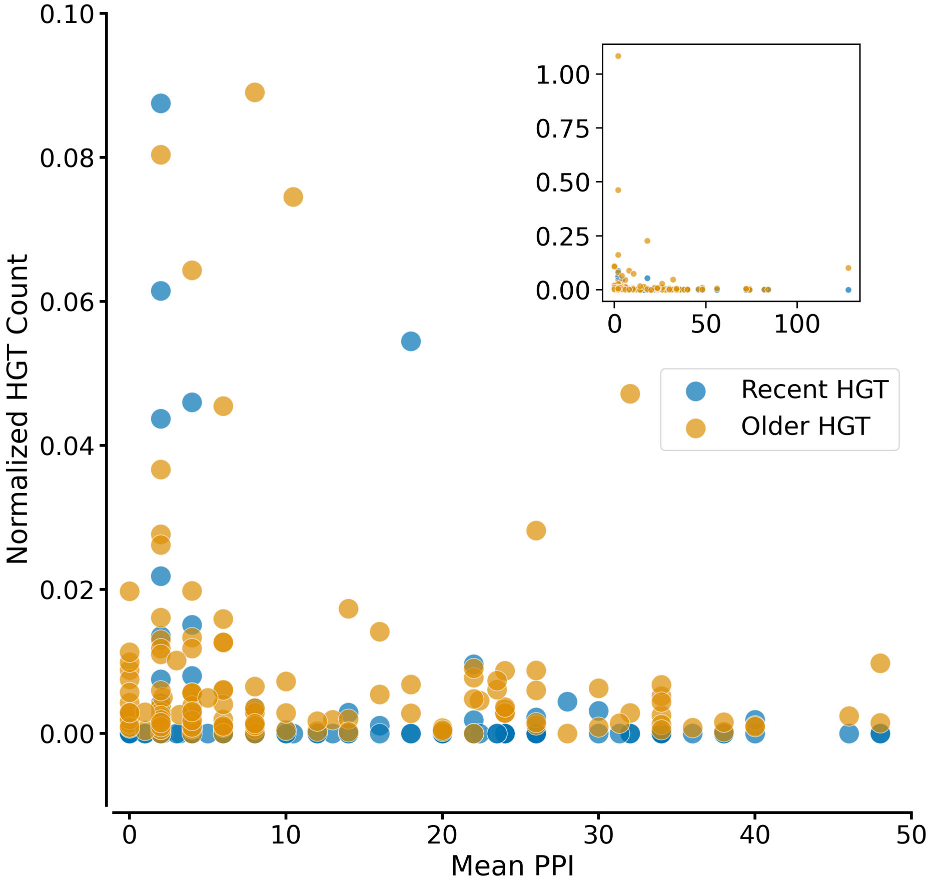


**Fig. S5. HGT counts correlate at most weakly with the number of protein-protein interactions.** Scatter plot of the normalized number of HGTs inferred for each gene family and its mean number of PPI (connectivity). The x-axis shows the mean PPI across all transferred genes in the gene family, and the y-axis shows the number of HGTs inferred for the gene family, divided by the total number of genes in the gene family for normalization. The inset shows the full figure, while the main figure shows a zoomed-in view of the data.
